# Gaze-Related Activity in Primate Frontal Cortex Predicts and Mitigates Spatial Uncertainty

**DOI:** 10.1101/2021.06.20.449147

**Authors:** Vishal Bharmauria, Adrian Schütz, Parisa Abedi Khoozani, Xiaogang Yan, Hongying Wang, Frank Bremmer, J. Douglas Crawford

## Abstract

A remarkable feature of primate behavior is the ability to predict future events based on past experience and current sensory cues. To understand how the brain plans movements in the presence of *unstable* cues, we recorded gaze-related activity in the frontal cortex of two monkeys engaged in a quasi-predictable cue-conflict task. Animals were trained to look toward remembered visual targets in the presence of a landmark that shifted with fixed amplitude but randomized direction. As simulated by a probabilistic model based on known physiology/behavior, gaze end points assumed a circular distribution around the target, mirroring the possible directions of the landmark shift. This predictive strategy was reflected in frontal cortex activity (especially supplementary eye fields), which anticipated future gaze distributions *before* the actual landmark shift. In general, these results implicate prefrontal cortex in the predictive integration of environmental cues and their learned statistical properties to mitigate spatial uncertainty.

## INTRODUCTION

A major purpose of the brain is to create predictive internal models of the surrounding environment to prepare for imminent action ^1,2^. This is challenging in a dynamic visual environment, with varying degrees of stability. But often we create expectations based on past probabilities, and these expectations manifest as behavioral strategies. For example, a soccer forward must integrate dynamic sensory information (goalie position relative to goal posts) with past knowledge of goalie behavior to aim the winning kick. Here, the forward is not just using visual landmarks to stabilize current spatial cognition ^3–9^, but also to generate predictions. The challenge here is that one of these landmarks (the goalie) is himself moving and only partially predictable. To mitigate this spatial uncertainty, some neural mechanism must integrate current sensory information with past experience.

The prospective influence of visual landmarks for predictive behavior has received little attention compared with their retrospective influence on spatial coding. For example, humans and non-human primates appear to optimally weigh allocentric and egocentric visual cues in cue-conflict tasks, e.g., where a shift in allocentric landmarks causes reach and gaze to deviate in the same direction ^6,10,11^. This behavior appears to involve neural computations in frontal cortex. In the absence of a visual landmark, gaze-related frontal activity simply grows more ‘noisy’ through time ^12–14^. However, in the presence of a shifting landmark, both the frontal (FEF) and supplementary (SEF) eye fields detect these shifts, ultimately integrating this information into their egocentric (eye-centered) gaze commands ^15,16^. However, other oculomotor studies suggest that these areas, especially the SEF, are involved in predictive gaze behaviors ^17–19^. We therefore hypothesized that frontal cortex (in particular SEF) might also be involved in predictive gaze behavior based on probabilistic spatial relations of environmental cues to future events.

We tested this hypothesis by simultaneously recording FEF and SEF neurons using the cue-conflict memory-guided saccade task developed and employed in our previous studies on the same animals ^11,15,16^ (**Fig. 1A**). In these previous studies, we showed a retrospective influence of a shifted visual landmark on gaze responses to a remembered visual target. But here, we focused on prospective coding, i.e., neural responses *before* the landmark shift. Guided by a theoretical framework based on prediction of probabilistic events and the neural computations noted above, we hypothesized that if the landmark shifted with a fixed amplitude but *random* direction, 1) animals might unconsciously develop a predictive gaze strategy to mitigate the future landmark influence ^1,2^, and 2) this strategy might be encoded *prospectively* in frontal cortex activity, particularly the SEF. Indeed, we found that, 1) animals developed a circular distribution of final gaze positions around the target, slightly biased toward the actual shift, and 2) both FEF and (especially) SEF neurons predicted these final gaze distributions just *before* the actual landmark shift. Collectively, these results implicate a critical role of frontal cortex in the integration of environmental cues and their learned statistical properties to predict and mitigate spatial uncertainty.

**Fig. 1.**
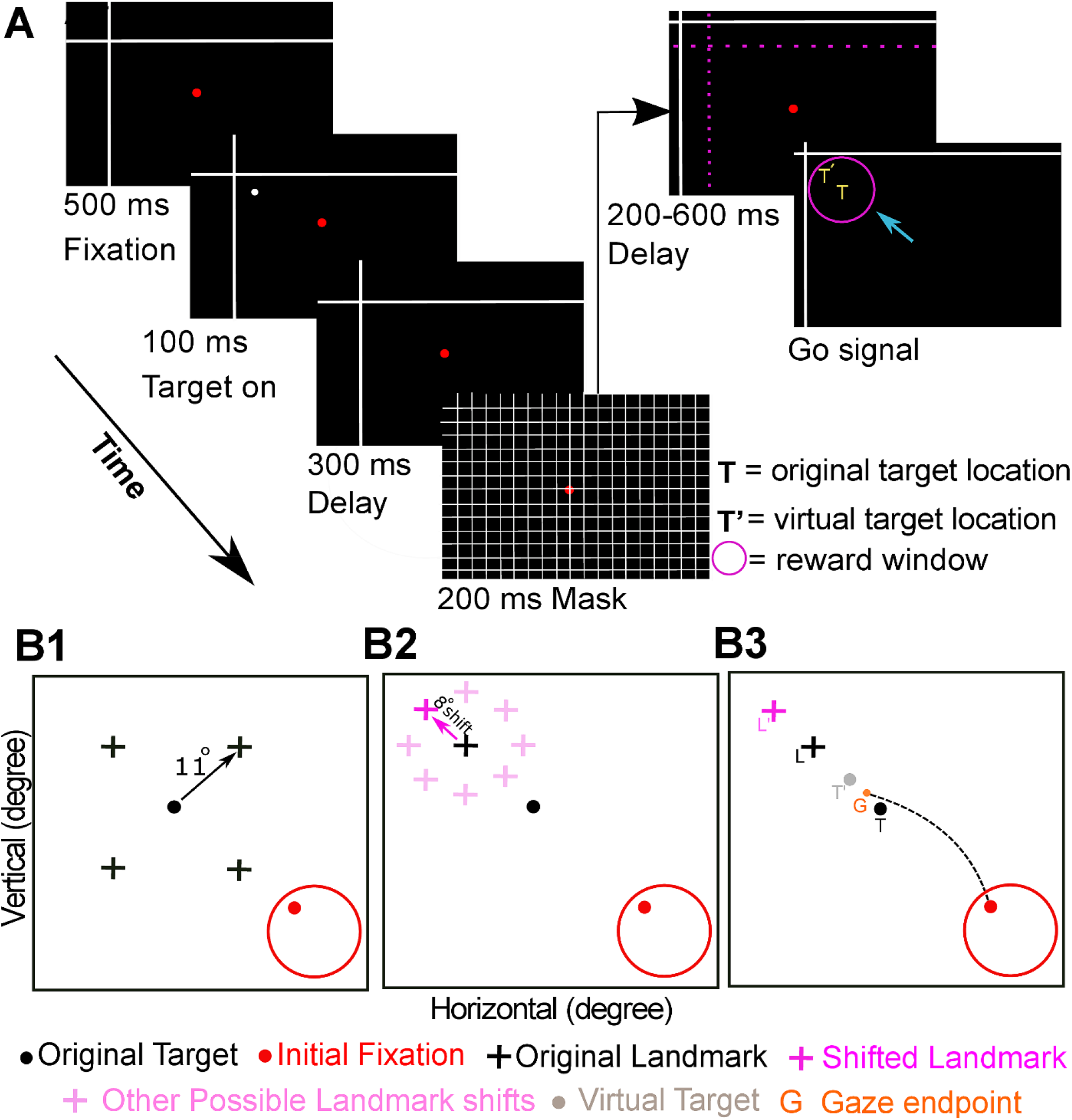
Experimental paradigm and behavior. **(A)** Cue-conflict experiment and its time course. The trial began by the monkey fixating on a red dot for 500 ms in the presence of a landmark (L, white intersecting lines) that was already present on the screen. A target (white dot) was then presented for 100 ms, followed by a first delay period of 300 ms and a grid-like mask (200 ms). After the mask, the landmark shifted (L’) in one of eight radial directions around the original landmark. Post-mask, and after a second variable memory delay (200-600 ms), the animal was cued (fixation dot off, i.e., go signal) to saccade to the remembered location of the target T. Accordingly, the animal was rewarded for landing its gaze (G) within a radius of 8-12° centered on the original target (i.e., either for looking at T = original target, at T’ = virtually shifted target fixed to landmark, or between T and T’). The cyan arrow denotes the head-unrestrained gaze saccade to the remembered location. Note for clarity purpose, the landmark shift is exaggerated in the figure. Importantly, the pink, yellow and cyan items were never present on the screen and are only shown for illustrative purpose. **(B1)** Schematic of four possible oblique landmark locations (black cross) in relation to a specific target (black dot). The red dot represents the initial eye fixation and the red circle corresponds to the typical fixation jitter. **(B2)** Schematic of a possible post-mask landmark shift (eight possible directions, 8° each, light pink) for an example shift (dark pink) away from the target. Note the radial distribution of possible landmark shifts around the original landmark. **(B3)** Schematic of a gaze shift (broken black line) with the gaze endpoint (G) between T and T’.

## RESULTS

### Task

To investigate how the brain might use visual landmarks to generate predictive gaze behavior, animals were trained on a cue-conflict task: a large landmark appeared in the background, then a target flashed briefly, followed by a surreptitious landmark shift (during a visual mask). Finally, animals were cued to aim their gaze toward the remembered target location **(Fig. 1A). Figure 1B** schematically shows 4 possible initial target-landmark configurations (**B1**) and the possible landmark shifts (**B2**). These shifts occurred in 1 of 8 directions around the original landmark location, but always had the same 8° amplitude, thereby forming a circular distribution. Animals were rewarded if gaze end points landed within 8-12° of the original target location (T, right panel), so that training did not bias their gaze behavior toward or away from the landmark shift. During experiments, the target position was varied throughout the visual field while randomly varying the relative landmark configuration and the direction of landmark shift. In previous experiments, we studied the influence of this landmark shift on subsequent premotor activity, and showed that it causes the one-dimensional distributions of final gaze position to shift in the same direction (**B3)** ^11,15,16^.

It is noteworthy that animals spent several months learning and performing this task for a water reward (see methods), so they had ample opportunity to implicitly learn its probabilistic properties (i.e., a fixed amplitude, variable direction landmark shift ^15,16^). To determine if these rules were incorporated into some predictive gaze control mechanism, here we analyzed final two-dimensional gaze distributions and examined neural activity *before* the actual landmark shift.

### Predictive Gaze Behavior: Actual and Simulated Distributions

#### Gaze Behavior

**Figure 2A** summarizes the distributions of gaze end points, for our two animals. Gaze distributions (blue-yellow color scale) are plotted relative to the remembered target position T (0°,0°; black dot), and the pink dots represent idealized target locations (T’) if they remained fixed to the landmark after shifting in the 8 possible directions (the dotted line connecting them represents the area where final gaze position gaze would result in a reward). The highest gaze densities (yellow) appear to cluster around the pink dots. At first glance one might assume that the animals simply waited for the landmark shift, and then deviated gaze in that direction, but in these plots, one cannot tell if there was any correlation between gaze and the actual shift direction.

**Fig. 2.**
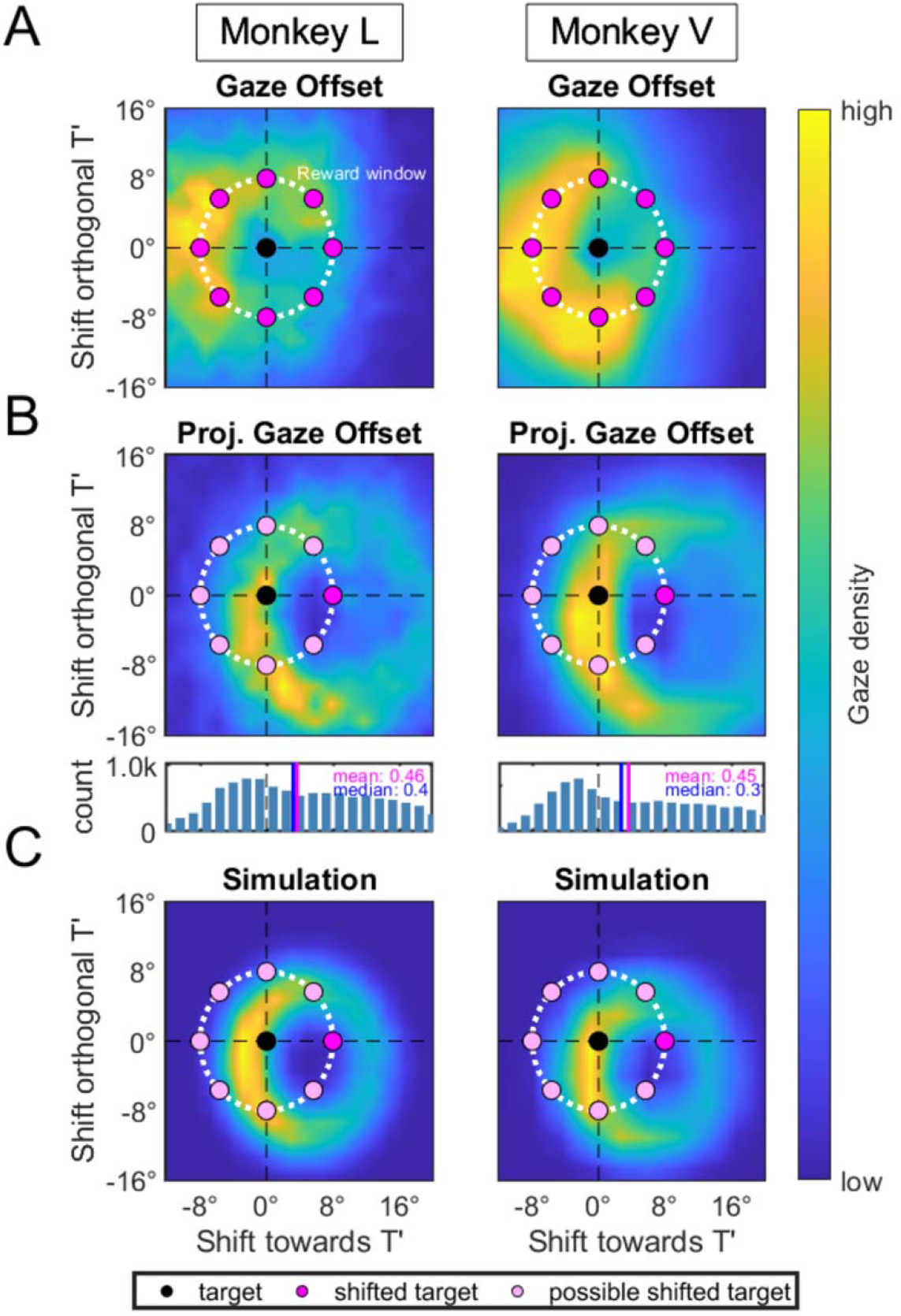
Gaze behavioral data and simulation. **(A):** 2-D distribution of the gaze endpoints (36084 trials in animal L, 27651 in animal V) relative to the actual location and directions of the landmark shifts. The black circle indicates the target position T, the pink circles indicate the shifted target positions T’. The x-component is given by the projection of the gaze endpoint scatter in the direction of the landmark shift and the y-component is given by the projection of the scatter orthogonal to the landmark shift. Note that these behavioral data were derived from the exact same trials used in the neurophysiological analysis provided below. (**B)** Displayed (top) is the normalized 2D distribution of the gaze endpoints shown in (A) around the target. All the shifted landmark positions (light pink circles) were graphically rotated such that they were all located to the right (dark pink circle). In other words, the light pink circles indicate shifted targets associated with landmark shifts that did not occur and the white dashed circle indicates the minimal reward window used in the experiment. The color map is indicative of the number of gaze endpoints in this region ranging from low (blue) to high (yellow). Gaze endpoints scatter in circular distribution with the highest density of gaze endpoints in a crescent area next to the target. Bottom: the 1D projection of the 2-D distribution of gaze endpoints along the direction of the landmark shift for both animals shows a bias in this direction. The blue line indicates the mean whereas the magenta line indicates the median. A comparison with the simulated data **(C)** shows the similarity between the real and simulated data for both monkeys.

To understand the real relationship between 2D gaze and the actual landmark shift, we rotated all of the data such that the direction of the actual landmark shift is always to the right (**Fig. 2B**). Now, the pink dot to the right represents the idealized target (T’), and the other 7 lighter dots represent the potential targets for the seven landmark shifts that did not occur. Gaze endpoints still produced a circular distribution (**Fig. 2B**, *upper panels*), reminiscent of the potential directions of the landmark shift. This pattern was also observed when each of the eight individual shift directions were analyzed separately (**Supplementary Fig. 1**). In other words, animals ‘guessed’ at the radial distribution of future landmark shifts, regardless of the actual direction of the landmark shift.

This does not mean that the actual landmark shift did not have an influence on gaze behavior. When data were collapsed into one dimension, i.e., shifts connecting T and T’ (**Fig. 2B, bottom panels**), they confirmed our previous findings ^11,15,16^: the overall gaze distributions were in fact shifted in the direction of the landmark shift (p < 0.01; Wilcoxon Rank Sum test), by a median of 3.2° in animal L and 2.4° in animal V. Thus, overall, both animals produced a predictive, circular distribution of gaze end points (similar to the possible landmark shifts) that was biased in the direction of the actual landmark shift.

#### Model

The behavioral data described above appears to support our hypothesis that animals learned to expect a fixed-amplitude landmark shift of varying direction. To understand how they might do this (and to make neurophysiological predictions), we developed a probabilistic model based on two known properties of the gaze control system, and one hypothetical property (see methods for mathematical description). The known properties are that 1) target memory is initially fairly precise but then progressively degrades through time, resulting in a broader distribution of variable gaze errors ^12,13,20^ and that 2) the landmark shift influences subsequent premotor codes, resulting in a shifted distribution of gaze end points ^15,16^. The third and novel component of the model is a ‘guess’ concerning the future landmark shift. Since the direction is unknown, this component results in a circular distribution of gaze estimates. We allowed these three model components (prediction, noise, actual shift influence) to “guess” a saccade vector and then calculated the weighted average across them to simulate the expected gaze distribution in our task.

After adjusting the model parameters (see methods), the simulated output almost exactly replicated the data (**Fig. 2C**), i.e., a ring-like distribution of gaze endpoints that was densest near the target but shifted in the direction of the landmark shift. (with a correlation of 0.81 and 0.81 between the actual and simulated data for monkey L and V respectively). Conversely, if we removed the predictive element of the model it resulted in a shifted gaussian distribution of gaze endpoints. Consistent with this, when we subtracted no-shift trials from the shift trials (**Supplementary Fig. 2**), the circular distribution collapsed to a shifted gaussian. These two findings confirm that the actual gaze distributions were a result of a probabilistic process, where landmark prediction explained the circular distribution, actual landmark influence explained the overall bias in this distribution, and interactions with a degraded target representation caused greater gaze density near the target. Again, the physiological basis of the latter two phenomena have already been described ^13,15,16^, but, the model makes a new and strong prediction: there must be some neural mechanism that predicts the future landmark influence before it actually happens.

### Neural Analysis: SEF predicts the future gaze distribution

The model described above suggests that the gaze control system implicitly anticipates the amplitude and guesses the direction of an impending probabilistic landmark shift, ultimately influencing the actual distribution of future gaze saccades. Based on the literature of oculomotor prediction ^21–24^, we expected the prefrontal gaze system, especially the SEF, to play a prominent role in this strategy. If so, their predictive neural signals should pass two criteria: 1) they should be present before the actual landmark shift, and 2) since the predictive strategy dominated final gaze position **(Fig. 2B)** then these signals should encode the observed deviations of final gaze from the original target.

To test this hypothesis, we analyzed early (pre-landmark shift) activity from 312 FEF and 256 SEF neurons recorded during the task described above (**Fig. 3A**). During experiments, we recorded neural response fields (the area of space that modulates neural activity). Targets were presented throughout the response field of each neuron, while randomly varying the 4 landmark configurations, and the 8 landmark shift directions. Consistent with previous studies, many of our neurons, especially in SEF ^25,26^, did not show significant spatial tuning. After removing these and applying our other exclusion criteria (see METHODS), we were left with 147 FEF and 68 SEF neurons for analysis. Mean spike density plots for these neuron populations, up until the landmark shift, are shown in **Figure 3B**. In both areas, visual targets evoked a strong visual response, followed by a lower-level memory response that lasted past the landmark shift ^15,16^. The 7 half-overlapping time steps shown above these plots show the temporal windows that we used in the following analysis. We then tested if activity predicted gaze in any of these periods.

**Figure 3.**
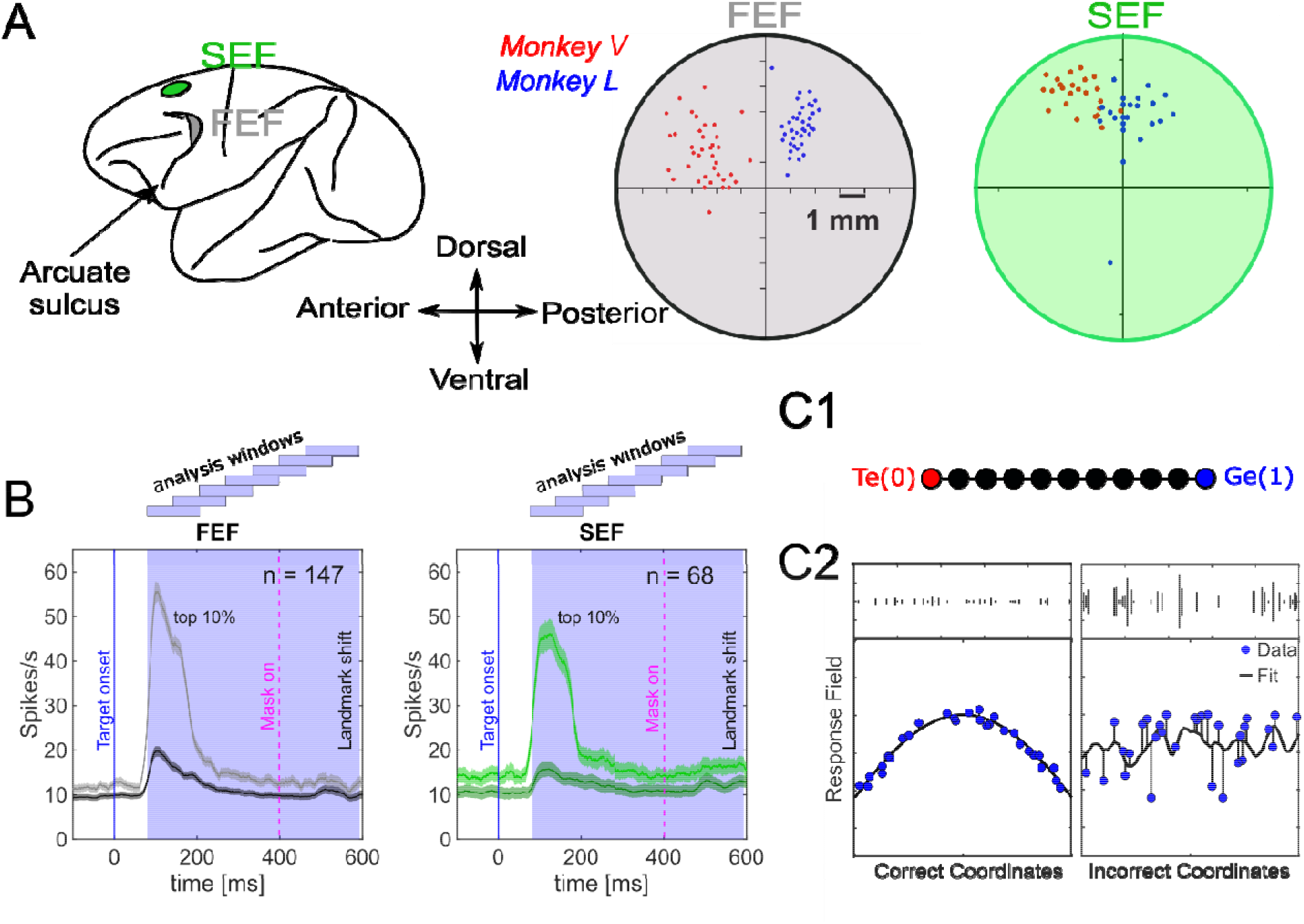
SEF and FEF recordings. **(A)** Left: The green and the gray sites represent the location of the SEF and FEF respectively. Right: Zoomed-in overlapped sections for FEF and SEF with sites of neural recordings (dots) that were confirmed with 50 µA current micro-stimulation. Blue and red dots correspond to recordings sites in Monkey L and Monkey V respectively. **(B)** Mean (± 95 % confidence) of the spike-density plots from target onset until landmark shift [dark; all trials from all neurons; light: top 10% best trials most likely depicting the hot spot activity of every neuron’s response field (RF) in visual responses, aligned to target onset (blue vertical line)]. The blue shaded region corresponds to the analysis window divided into 7 half-overlapping x ms wide time-steps, as depicted above the shaded area. **(C)** A schematic behind the logic of response field analysis. **(C1**) Shown is a schematic of the continuum between Te(0) and Ge(1) with intermediate steps. **(C2**) The X-axis denotes the coordinate frame, and the Y-axis represents the corresponding activity. Briefly, if the activity related to a specific target is plotted in the correct/best reference frame, this will result in lowest residuals, i.e., if the neural activity to a target is fixed (left) then the data (blue) would fit (black curve) better on that, yielding lower residuals compared with when the activity is plotted in an incorrect frame, yielding higher residuals (right).

To do this, we characterized if neurons were coding original target location (T), the future final gaze position (G), or something in between, called the ‘T-G continuum’ (discretized in ten steps), calculated relative to initial eye orientation ^12,13,15,16,27^. In this analysis, a value of 0 indicates a pure target-relative-to-eye encoding, while a value of 1 indicates a final-gaze-relative-to-eye encoding, values between 0-1 indicate an intermediate code, and values beyond 0 / 1 could indicate a negative (perhaps inhibitory) influence of the opposite factor (**Fig. 3C1**). This analysis allowed us to plot the response field data in each of these coordinate frames, and to perform a non-parametric fit to each dataset ^28^. The one spatial step (out of ten, see above) that yielded the lowest residuals (deviations) between the actual neural responses and the fit was deemed to provide the best fit and hence indicate the coordinate system employed by a given neuron at a given time (**Fig. 3C2)**. We performed this analysis for each of the time steps shown in **Figure 3B**, to track the temporal evolution of the spatial coding before the landmark shift. Note since G is derived from the *actual* gaze data constituting the predictive distribution in **Figure 2**, neurons / populations that approach G must be involved in prediction.

**Figure 4** shows a typical example of the response field fitting for an FEF (upper row) and an SEF (lower row) neuron. The leftward panels (A, D) display raster with the spike density plot for these neurons, aligned to the target onset (blue arrow). The blue shaded area corresponds to the analysis window that is divided into our 7 half-overlapping time-steps. To the right of these plots are response fields calculated at the 2^nd^ and 5^th^ time steps (indicated by green rectangles in A/D) and plotted in their best T-G coordinate frame (indicated by the yellow dot on the scale above each plot). Each circle in the response field map corresponds to neural activity from a single trial, where the larger the circle the larger the response (i.e., number of action potentials). The colored heat maps represent the non-parametric fit to these data, where red depicts the ‘hot-spot’.

**Figure 4.**
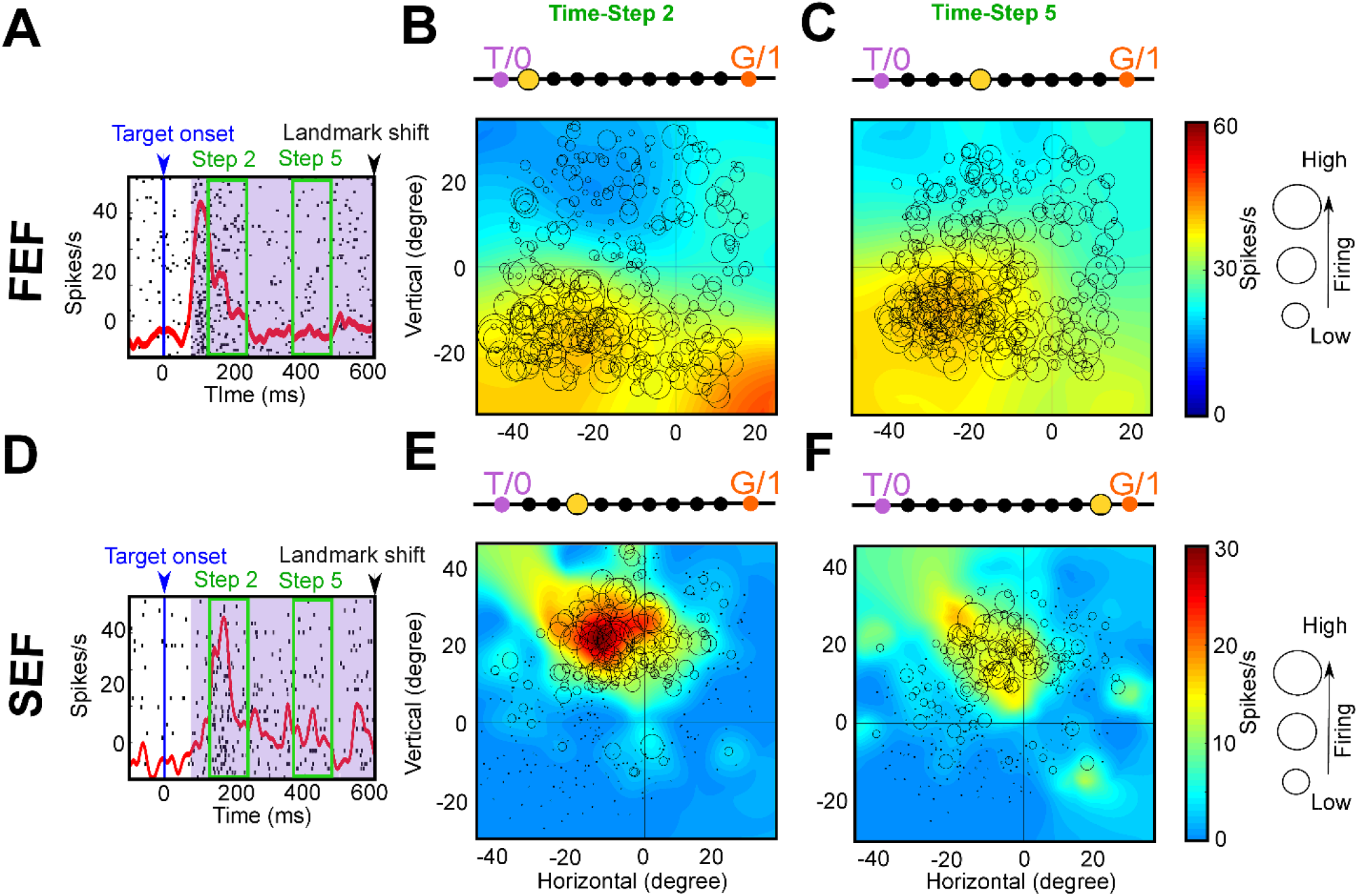
Typical examples of spatial encoding of an SEF and FEF neuron. **(A)** Raster with spike density plot (red curve) for the FEF neuron. The blue arrow corresponds to the target onset and the blue shaded area represents the analysis window divided into 7 half-overlapping time-steps. The green rectangles correspond to the time-steps 2 and 5. **(B)** Response field plot at time-step 2. The response field fits at 1^st^ point from T. The yellow blob represents the hot spot of the response field. **(C)** Response field plot at time step 5 and it fits best at 4^th^ step from T. **(D)** Same convention as A but for SEF neuron. **(E)** Response field plot at time-step 2 and it fits best at 3^rd^ point from T. **(F)** Response field at time step 5 and it fits best at 9^th^ point from T suggesting a predictive shift toward gaze. The color bar stands for both response fields. The circle size is proportional to response magnitude. Note: 0,0 denotes the center of the coordinate system (the fovea) that yielded lowest residuals (best fit).

At time step 2 (**Fig. 4 B/E**; spanning the late phasic response to target presentation), both the SEF and FEF examples show a best fit near T, indicating that these neurons were coding target location relative to the eye. At time step 5 (**Fig. 4 C/F**; just after mask onset, and just before the anticipated landmark shift), there were no obvious shifts in the response fields. However, there were shifts in the best T-G fits, signifying a change in the underlying neural code. In the FEF example, there was a 30% shift toward G, signifying a closer relation to future gaze position. Further, the SEF example shifted 90% toward G. This means that this SEF neuron was predicting the circular distribution of gaze deviations from T, on a trial-by-trial basis, just before the actual landmark shift.

To document these observations through time, we pooled the T-G fits across all FEF (n =147) and across all SEF (n = 68) neurons and then analyzed each population code as a function of time (**Fig. 5**). **Figure 5A** illustrates the mean spike density plots for the SEF and FEF neurons across 7 time steps ranging from visual response onset until the (invisible) landmark shift. The pink shaded area corresponds to the duration of the mask.

**Figure 5:**
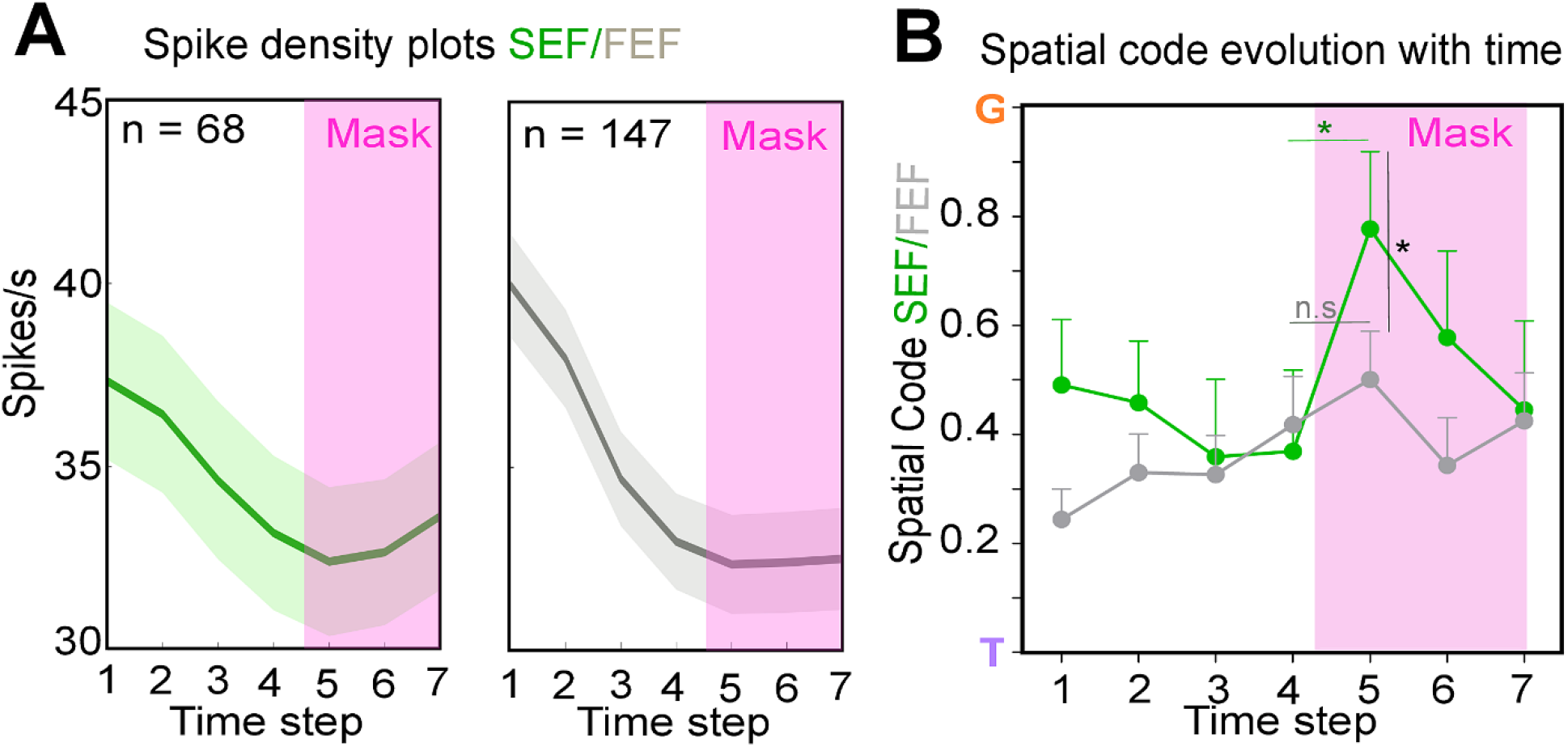
Spatial code evolution with time. **(A)** Spike density plots (mean ± SEM) for SEF (green) and FEF (gray) from visual response onset until the landmark shift divided into 7-half overlapping time-steps. **(B)** Spatial code evolution with time for SEF (green) and FEF (gray) neurons along the target-to-gaze (T-G) continuum. A sudden predictive shift toward G was noticed for SEF neurons at 5^th^ step that significantly differed from corresponding FEF step (p = 0.028, unpaired t-test) and the 4^th^ SEF step (p = 0.02, unpaired t-test).The pink area corresponds to the duration of the mask.

Both the FEF and SEF showed significant deviations from T at all time steps, which could partially be accounted for by the degraded T representation in our model. However, they followed different time courses. For the FEF (grey symbols and curve), there was a gradual progression from T towards G coding along the time-steps, as noted previously ^12,15^. However, for the SEF population (**Fig. 5B**, green symbols and curve) the spatial code already started midway between T and G at time-step 1 (the visual response to the target in the presence of the landmark) with a significantly greater shift than FEF (p=0.01, Mann-Whitney test). Then, after reverting toward the FEF curve for several steps in the memory period, the SEF again displayed a sudden shift of 78 % toward G at time-step 5 (just after mask onset and just before the probabilistic landmark shift), with a significantly greater shift than FEF (p = 0.028, Mann-Whitney test). Further, there was a significant difference between the 4^th^ and 5^th^ steps for the SEF (p = 0.02, unpaired t-test). The FEF appears to follow a small trend at this point, but this did not reach significance (p > 0.05, Mann Whitney test) population. These data suggest that the SEF played a special role in predicting future gaze direction, just before the landmark shift, including (and possibly causing) the trial-to-trial ‘guess’ at the direction of landmark shift.

To illustrate how this predictive shift occurred across the full distribution of our FEF and SEF populations, we computed ‘violin’ plot fits to the T-G distributions of all spatially tuned neurons **(Fig. 6)** for time steps 1-5 (where the G prediction peaked). These populations showed means and medians between T and G, but extend beyond T and G, a phenomenon that has been noted in previous studies of intermediate reference frames in both real and artificial neural populations ^12,15,16,20,29–31^. Both populations revealed relatively stable code distributions for the first 3 time steps. The FEF showed a simpler distribution that remained fairly stable, except for the slight expansion of a bimodal ‘head’ at time steps 4 and 5. In contrast, the more complex SEF population distribution started to shift upward at step 4, with a dramatic upward shift at step 5.

**Figure 6:**
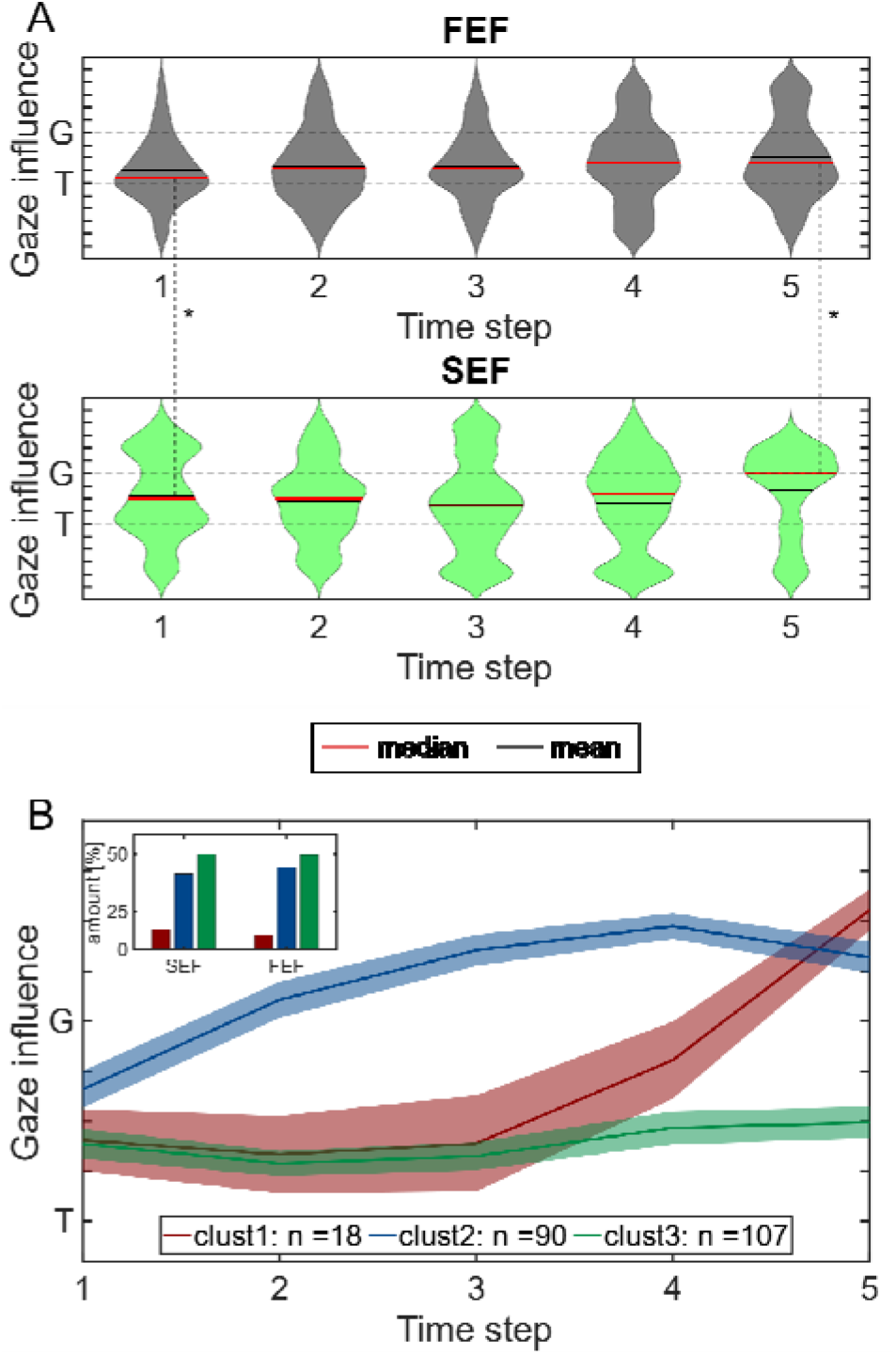
Detailed evolution of spatial codes through time. **(A)** Violin plots for FEF (top) and SEF (bottom). The width of each plot indicates the relative number of neurons with fits at a particular point on the T-G continuum. The mean of each distribution is indicated by the red line and median by the blue line. Such plots combine the strengths of bar graphs and frequency histograms (arranged in the vertical dimension), **(B)** Dissociation of neurons into three distinct (colour coded) clusters of neurons using a clustering approach. The main graph plots means and confidence intervals of T-G fits for the combined FEF/SEF population, plotted through five time steps. Inset shows the relative numbers of neurons in SEF and FEF that fit within these three clusters.

Finally, to test if distinct sub-populations of neurons contribute differently to these coding shifts through time, we employed a dimensionality reduction approach on the first five time-steps by hierarchical clustering. To be objective, (and because FEF and SEF are highly interconnected with similar responses) we pooled neurons from both areas for this analysis. We then used the Ward method in conjunction with the Euclidian metric (see methods) to identify clusters of neurons within the spatiotemporal (T-G fit versus time) coding patterns of the entire population. This resulted in three distinct neuron clusters (**Fig. 6B)**, somewhat reminiscent of the three components in our model. Cluster 1 (red) neurons showed a predictive shift toward G beginning at step 3 and peaking at step 5, resembling the predictive response seen in the whole population analysis. Cluster 2 (blue) neurons reached and maintained preference gaze coding as early as the second step, whereas cluster 3 (green) neurons maintaining a slightly degraded target code. Proportionately, more SEF neurons (10.3%) participated in cluster 1 compared to FEF (7.5%), whereas both areas contributed nearly equally (39.7% SEF/ 42.8% FEF in cluster 2; 50% SEF/ 49.7% FEF in cluster 3) to the other clusters (**Fig. 6B**, *inset*). This analysis suggests a considerable degree of signal sharing between FEF and SEF, but this signal distribution manifests itself differently in their whole population codes (**Figs. 5, 6A**). This sharing may explain why the overall FEF population shows a small trend toward gaze prediction, whereas the SEF explicitly predicts final gaze direction, coding (and perhaps producing) a strategy to mitigate the expected future landmark influence.

## DISCUSSION

To investigate how the frontal cortex (FEF and SEF) integrates environmental cues and learned probabilities for predictive gaze behavior, we used a cue-conflict memory-guided saccade task, where a visual landmark shifted in a quasi-predictive radial pattern after a mask. We found that: 1) final gaze formed a circular pattern around the original target, resembling the shift probability distribution but slightly biased in the direction of actual shift, 2) a probabilistic model of the above data yielded a circular pattern that was strikingly similar to the real data. 3) this behavioral strategy was reflected in supplementary eye field response fields, which showed a transition to gaze coding just before the actual landmark shift and 4) a clustering algorithm dissociated three types of neurons in both areas, suggesting a shared modular specificity. Collectively, this study provides new insights into how the brain uses visual cues for predictive, probabilistic gaze behavior, especially in a dynamic but quasi-predictable visual environment.

### Relation to previous behavioral studies

Various previous studies have addressed the use of landmarks in the retrospective coding of target memory for action planning ^32–35^, and other studies have considered the prospective use of cues for predictive gaze coding ^2,36,37^, but here have we considered the combination of these two factors for spatial behavior involving probabilistic environmental cues. In our task, an environmental cue that would normally augment visual stability ^38,39^ becomes unstable. Imagine if you used a certain landmark to navigate to work every day, but some malicious prankster started relocating it every night. After a while, one might learn to predict and mitigate the effects of this trick, either by choosing other landmarks, or learning the trickster’s pattern. Although our task was visually impoverished compared with this example, the general principle of combining environmental cues and prediction based on prior knowledge appears to be a central (some might say primary) aspect of gaze control and brain function in general for real world behavior ^37,40^. Thus, although the mechanisms observed here pertain to a very specific task, they likely generalize to many other daily tasks, i.e., wherever there is spatial uncertainty in our future environment.

In the gaze control system, it has been suggested that spatial predictions based on environmental cues guide goal selection ^36,37,40,41^. Prior knowledge/memory representation facilitates visual search ^42^, influences goal-directed movements to the target ^43,44^, allows predictions based on the history and motion of a target ^45–47^. Moreover, it has been proposed that many aspects of behavior are governed by Bayesian models. Previous studies have shown that the brain integrates visual landmarks with target information in a Bayesian fashion for gaze control ^11,15,48,49^ and other goal-directed movements ^5,6,50,51^. In one study ^52^, a target acquisition model (TAM) based on a *target map* (essentially the proposed/possible locations for gaze in a defined scene) exhibited similar levels of performance as human participants for a target search from a set of previewed targets and identical display later on, suggesting that the brain creates a probabilistic map of possible targets. Furthermore, it is widely shown that the brain creates cognitive maps through repetitive reinforcement, learning, prediction and reward maximization ^53,54^.

The landmark shifts in this experiment were masked, but even if monkeys ‘noticed’ the change in position, it seems unlikely that they developed a ‘conscious’ predictive strategy to deal with the landmark shifts. For example, humans are influenced by landmark shifts even when told to ignore them ^6^. Instead, it seems likely that their strategy was learned implicitly over the course of many thousands of trials during training and data collection. The constant repetition of a landmark shift with a fixed amplitude but variable direction may have allowed the brain to generate a probabilistic map of the distribution of possible landmark shifts, as in our model. And the actual influence of a landmark shift appears to be developed naturally as a prior ^2,37,52^. By combining a probabilistic map with a noisy gaze distribution and the influence of the actual target shift, our model was able to replicate the actual gaze strategy (**Fig. 2 C**). But how could the brain achieve this?

### A neural algorithm for landmark-based gaze prediction

In this section we link the behavioral data to our neurophysiology by speculating how the steps in our model could relate to internal brain events. In **Figure 7**, we have speculatively superimposed simulations of the three main model components (*Rows R1-3*) against the seven main events of our task (*Columns C1-7*). Each panel represents the contribution of the corresponding model component to the relevant event. As in **Figure 2C**, the simulation shows probability distribution of gaze end points around the target, superimposed on a circle showing the possible directions of the landmark shift (with the actual shift direction normalized to the right). *Row 1* illustrates a Gaussian representation of target position, which is initially fairly precise (*R1,C2*) but then progressively degrades through time, resulting in a broader distribution of variable gaze errors by the time of the final gaze command (*R1,C6*). This has already been observed both in behavior and in FEF memory responses ^12,13^. It is noteworthy that in our model this area corresponded to the spatial ‘reward window’ provided to the monkey, suggesting a constrain related to reward maximization ^55^. *Row 3* shows the influence of the actual landmark shift, resulting in a partial shift in the gaze distribution in the same direction (*R3,C4*). This has been observed in the premotor FEF/SEF responses that follows the landmark shift ^15,16^ and can be explained by optimal integration theory ^6,56^.

**Figure 7:**
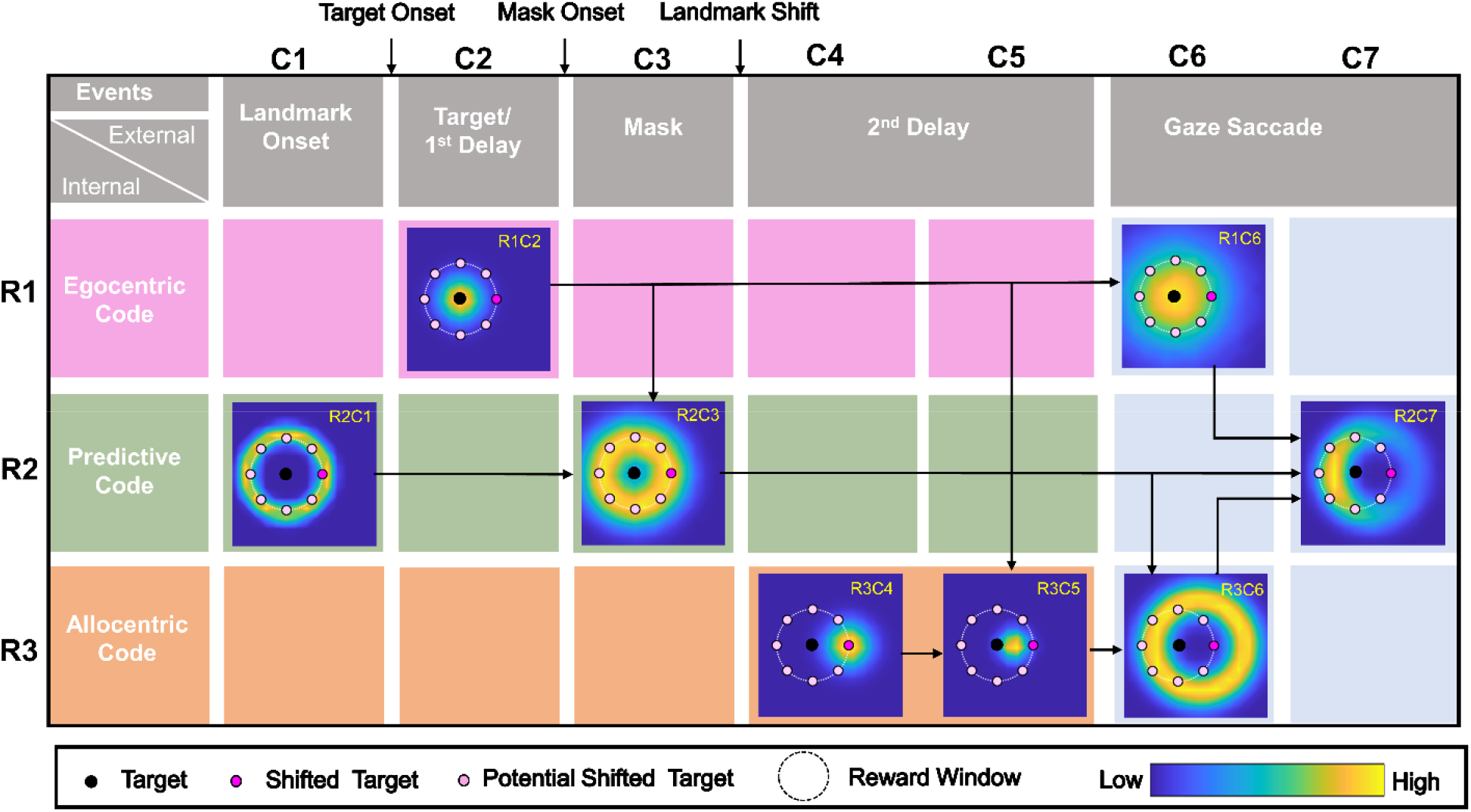
Schematic representation of the contributions of the three major components of our model (rows R1-3) with respect to the 7 major events in our task (Columns C1-7). The small pictograms show 2D simulations of gaze distributions produced by the model components during key events. As in **Figure 2C**, simulations are directionally normalized so that the landmark shift is to the right. In each simulation, the black dot represents the target, the magenta dot represents the virtually shifted target (T’), the light-colored magenta dots represent potential target shifts that did not occur, and the dashed white circle indicates the minimal reward window used during the experiment. See Results text for explanation and methods for mathematical details of the simulations.

*Importantly, row 2* shows the novel aspect of the model. Here, our SEF data suggest that predictive information about the future landmark influence is already present (perhaps at the synaptic level) when the visual target response interacts with the background landmark (*R2,C2*). Second, the high (78%) SEF gaze prediction during this mask (*R2,C3*) suggests that the mask might ‘warn’ of the upcoming landmark shift, triggering a comparison between the target representation and the landmark prediction that produces the future circular gaze distribution (minus only the bias due to the actual shift). Finally, when this probability distribution combines with the other two probability distributions (gaussian gaze error and influence of the landmark shift) to produce the final motor command (*Column 7*), it results in a ring-like distribution of gaze end points that is somewhat denser near the target but shifted in the direction of the landmark shift. Note that individual trials are directed pseudorandomly (as in our data) but the overall gaze distribution maximizes reward across trials. Overall, this strategy maximizes reward outcome based on visual cues and their link to expected probabilistic events ^57,58^. In lay terms, the model makes an educated ‘guess’. Accordingly, this approach provides a model framework for understanding how neurons might actually implement such algorithms.

### Neural Implementation: role of the SEF and FEF

While both the FEF and SEF showed a trend toward gaze coding early in the task, the slow rise in FEF could be interpreted as noise accumulation ^12,13,15^. However, the SEF passed both our criteria for predictive coding: it showed a sudden shift toward gaze coding (along with its prediction-dominated circular pattern of gaze deviations) just *before* the actual landmark shift. FEF and SEF are reciprocally interconnected and show similar properties, but the general consensus is that the FEF is more tightly linked to the generation of saccades. In contrast, the SEF holds ‘executive’ control and influences oculomotor centers with a multitude of signals such as reward, prediction, decision making, learning, rank dependency, surprise, conflict monitoring and behavioral supervision ^22,24^. Both areas are involved in eye-centered and allocentric visuomotor transformations ^12,13,15,16^, but the SEF is also implicated in object-centered coding ^59,60^. Furthermore, the SEF encodes two types of errors that are relevant for learning and prediction: 1) amount of reward, and 2) subjective probability of feedback ^21^.

Based on our model (**Fig. 7**) and the general principle of reward/effort maximization ^57,58^, we propose the following explanation for our neurophysiological data. Our previous results suggest that the FEF and SEF continue to show an eye-centered target-relative-to-eye to gaze-relative-to-eye transformation for saccades in the presence of a landmark ^15,16^, but their visual signals are influenced by landmarks in a fashion that depends on target-landmark configuration ^32^. Thus target-landmark configuration information is present from the start of each trial, but our new data here suggest that these interactions are influenced over time by the expectation of future probabilistic events and reward ^16,61^.

Overall, the three population clusters identified in **Figure 6** are somewhat reminiscent of the three conceptual channels in our model, but the analogy is not perfect. One (**Fig. 6B:** green) seems to maintain a slightly noisy target code, one (red) appears to be involved in prediction just before the landmark shift, and the progressive transition in the third (blue) could be interpreted either as noise build up or prediction. However, one cannot know if the analysis algorithm is separating clusters on the same basis as our conceptual model.

Despite the general similarities between FEF and SEF distribution across clusters, our current data suggest that only the SEF plays a stronger role in the predictive gaze strategy: only SEF shows a significant shift toward gaze coding just before the landmark shift, perhaps triggered by visual input from the mask. Although it is not possible to infer causality from neural activity alone (particularly in such a highly interconnected system), we propose that the circular distribution of gaze positions around the target originates in the predictive SEF code. Interestingly, this predictive peak in the SEF coding then dissipates somewhat, perhaps exerting its influence thereafter through synaptic modulation of distribution of signals across the SEF — posterior parietal cortex, dorsolateral prefrontal cortex — FEF memory loop ^62–64^. Most likely prefrontal predictive activity influences final motor output in both structures, because SEF and FEF motor responses encode future gaze position in this task ^15,16^, which must include the circular patterns observed here in the behavior. One would expect the same to hold true in the motor response of the superior colliculus.

As we observed previously, the actual landmark shift influence appears during the following delay activity in both the FEF and SEF, through slightly different and complementary mechanisms (specifically the balance of activity in visuomotor vs. motor neurons ^15,16^). This would implement the shift in the ‘donut’ shown here (**Fig. 2**). Initially these allocentric and egocentric signals were multiplexed in separate codes, but became fully integrated in the final motor response, as they must to influence the actual behavior.

In short, we are able to explain most of the behavior described here in terms of our own data and previous literature, with a minimum of speculation. This explanation is admittedly highly specific to the current task and training, but it is exceedingly unlikely that these circuits developed for such a specialized purpose. More likely, the circuits described here illustrate the flexible capacity of this system to contribute to predictive strategies based on learned environmental heuristics, and thus should generalize to other situations.

### General Conclusions and implications

Prediction is fundamental to brain function and gaze behavior ^37,65^, but becomes challenging when environmental cues themselves are unstable. In such situations, the brain can only incorporate experienced statistical properties of the environment, and then essentially ‘guess’ at the properties that remain uncertain. Using a quasi-predictable gaze paradigm involving a series of visual cues (a landmark, a target, a mask, and landmark shift in an unpredictable direction) we showed that 1) Rhesus macaques developed a predictive strategy to — most likely implicitly — anticipate the future consequences of a probabilistic landmark shift, and 2) that frontal cortex (SEF in particular) carries and perhaps produces the predictive signals that underlie this behavior. This shows that frontal cortex is involved in the use of environmental cues and the learned statistics of their future motion to generate predictive behaviours. It is likely that this role of frontal cortex generalizes to other visual behaviors, i.e., whenever movements are planned in the presence of spatial uncertainty. Conversely, frontal damage should adversely affect one’s ability to generate predictive behavior in a dynamic environment.

## MATERIALS AND METHODS

### Surgical Procedures and Recordings of 3D Gaze, Eye, and Head

The experimental protocols followed the guidelines of Canadian Council on Animal Care on the use of laboratory animals and were also approved by the York University Animal Care Committee. Neuronal recordings were done on two female *Macaca mulatta* monkeys (Monkey V and Monkey L). Their left eyes were implanted with 2D and 3D scleral search coils for eye-movement recordings ^66,67^. The eye coils permitted us to register 3D movements of the eyes (i.e., gaze) and orientation (horizontal, vertical, and torsional components of eye orientation relative to space). Two head coils (orthogonal to each other) were also connected during the experiment that allowed similar recordings of the orientation of the head in space. Then, in both animals a recording chamber was implanted on FEF and SEF, centered in stereotaxic coordinates at 25 mm anterior and 19 mm lateral for FEF and 25 mm anterior and 0 mm lateral for SEF. A craniotomy of 19 mm (diameter) on FEF and SEF covering the chamber bases (adhered over the trephination with dental acrylic) allowed access to the right FEF and SEF. Animals were seated within a custom-made primate chair during experiments, allowing free head movements at the center of three mutually orthogonal magnetic fields ^66^. The values recorded from the 2-D and 3-D eye and head coils allowed us to compute other variables such as eye orientation relative to the head, eye- and head-velocities, and accelerations ^66^.

### Basic Behavioral Paradigm

The visual stimuli were presented on a flat screen (placed 80 cm in front of the animal) using laser projections **(Fig. 1A)**. The animals were trained on a standard memory-guided saccade task where they had to remember a target location relative to a visual allocentric landmark (two intersecting lines). This led to a temporal delay between the presentation of the target and beginning of the eye movement. The experiment was conducted in dark to avoid any other allocentric cue. A single trial consisted of the animal fixating on a red dot (placed centrally) for 500 ms in the presence of the allocentric landmark. This was followed by a brief flash of the visual target (T, white dot) for 100 ms, and then a brief delay (300 ms), a grid-like mask (200 ms, this hides the past visual traces, and also the current and future landmark) and a second memory delay (200-600 ms, i.e., from the onset of the landmark until the go signal). As the red fixation dot extinguished, the animal was signaled to saccade head-unrestrained (indicated by the solid green arrow) toward the memorized location of the target either in the presence of a shifted landmark (90 % of trials) or in absence of it (10 %, no-shift/zero-shift condition, i.e., the landmark was present at the same location as before mask). These trials with zero-shift were used to compute data at the ‘origin’ of the coordinate system for the T-T’ spatial model fits as described below. The saccade targets were flashed one-by-one randomly throughout the response field of a neuron. Note: magenta color highlights the items that were not presented on the screen (they are shown only for representational purposes).

The spatial details of the task are depicted in **Figure 1B** illustrating the gaze shift (blue curve) to an example target (T) in presence of a shifted landmark (L’). **Figure 1B1** shows possible original landmark locations (L, black cross) to an example target (black dot). The red dot corresponds to the eye fixation and the red circle represents the jitter in initial home fixations. The landmark vertex could initially appear at one of four locations, 11° obliquely relative to the target. **Figure 1B2** illustrates possible landmark shifts (magenta crosses) to an example original landmark location. In this case the landmark shifted *(8°)* to the top left as depicted by the black arrow. Notably, *the timing and amplitude of this shift was fixed*. **Figure 1B3** shows an example gaze shift from initial eye fixation to final gaze endpoint (G). T’ stands for the virtual target (fixed to the shifted landmark). Since these animals had been trained, tested behaviorally ^11^ and then retrained for this study over a period exceeding two years, it is reasonable to expect that they may have learned to anticipate the timing and the amount of influence of the landmark shift. However, we were careful not to bias this influence: animals were rewarded with a water-drop if gaze was placed (G) within 8-12° radius around the original target (i.e., they were rewarded if they looked at T, toward or away from T’, or anywhere in between). Based on our previous behavioral result in these animals ^11^, we expected this paradigm to cause gaze to shift partially toward the virtually shifted target in landmark coordinates (T’).

Note that this paradigm was optimized for our method for fitting spatial models to neural activity (see below), which is based on variable dissociations between measurable parameters such as target location and effectors (gaze, eye, head), and various egocentric / allocentric reference frames ^12,28^. This was optimized by providing variable landmark locations and shift directions, and the use of a large reward window to allow these shifts (and other endogenous factors) to influence gaze errors relative to T. We also jittered the initial fixation locations within a 7-12° window to dissociate gaze-centered and space-centered frames of reference (note that no correlation was observed between the initial gaze location and final gaze errors). Further dissociations between effectors and egocentric frames were provided by the animals themselves, i.e., in the naturally variable contributions of the eye and head to initial gaze position and the amplitude/direction of gaze shifts. Details of such behavior have been described in detail in our previous papers ^12,28^.

### Behavioral Recordings and Analysis

During experiments, we recorded the movement of eye and head orientations (in space) with a sampling rate of 1000 Hz. For the analysis of eye movement, the saccade onset (eye movement in space) was marked at the point in time when the gaze velocity exceeded 50°/s and the gaze offset was marked as the point in time when the velocity declined below 30°/s. The head movement was marked from the saccade onset till the time point at which the head velocity declined below 15°/s.

When the landmark shifted (90% of trials), its influence on measured future gaze position (G_i_) was called projected gaze offset (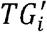), computed as follows:

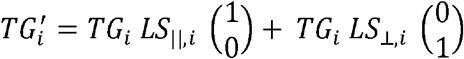

where 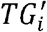 is allocentric weight; *LS*_‖,*i*_ is the landmark shift in trial i, *LS* _⊥,*i*_ is the landmark shift rotated by 90° counterclockwise and *TG*_*i*_ is the gaze offset (difference between the actual target location and the final measured gaze position). This computation was done for each trial i, and then averaged to find the representative landmark influence on behavior in a large number of trials. A projected gaze offset 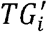 of 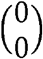 signifies a gaze shift that landed exactly on T. A projected gaze offset 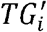 of 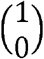 means that the gaze headed toward a virtual target position (T’) that remained fixed to the shifted landmark position. A projected gaze offset 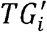 of 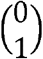 means that the gaze headed towards a point rotated by 90° counterclockwise virtual target position (T’).

### Simulation

We were interested in how the behavior data can be explained. To this end, we designed a stochastic process serving as means to simulate the neuronal process leading up to the behavior **(Fig. 7)**. The stochastic process starts with three base distributions. The three distributions represent the visual input in one of three codes, egocentric 𝒫_*ego*_(C2R1) and predictive codes 𝒫 _*pred*_(C1R2) and allocentric influence 𝒫_*allo*_(C4R3).

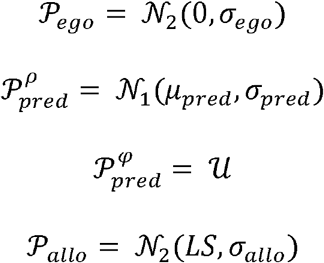

These distributions are then sampled resulting in three “guesses”:

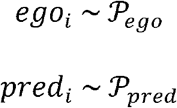

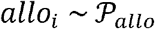

Then the weighted average of these guesses is calculated resulting in the intermediate distribution 𝒫 _*ego,pred*_ (C3R2), 𝒫_*ego,allo*_ (C5R3) and 𝒫_*ego,pred,allo*_(C6R3).

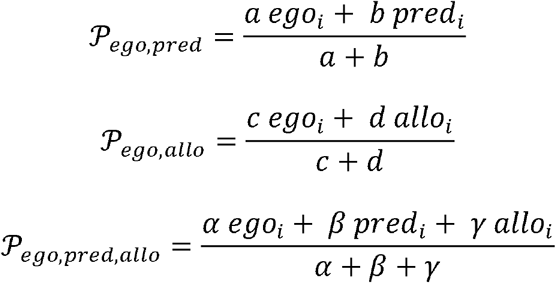

Finally the combined distribution is reweighted by a faded ego centric target memory

*𝒲*_*ego*_(C6R1) resulting in the final distribution 𝒫 (C7R2).

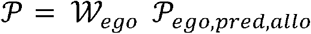

To produce one simulated saccade this distribution 𝒫 is sampled. This sampling was repeated 10000 times. The results of this process are displayed in **Figure 2**.

### Cluster analysis

To visualize the dynamics of coding of single units, we aimed to reduce the time course to a small number of archetypical time courses. This dimensionality reduction was achieved by hierarchical clustering. For the clustering, we considered the coding and tuning time courses of the individual neurons. We employed the ward method ^68^ in conjunction with the Euclidian metric. The clustering resulted in three distinct clusters representing archetypical single neuron time courses. The average time courses for each of these three clusters are shown in **Figure 6B**.

### Electrophysiological Recordings and Response Field Mapping

We lowered tungsten electrodes (0.2–2.0 MΩ impedance, FHC Inc.) into the FEF and SEF [using separate Narishige (MO-90) hydraulic micromanipulators for each area] to record the neuronal activity. We then digitized, amplified, filtered, and saved the recorded activity for offline spike sorting. Sorting was performed using template matching and the principal component analysis on the isolated clusters (done with Plexon MAP System). The recorded sites (in head-restrained conditions) were further confirmed by low-threshold electrical microstimulation (50 μA) ^69^. The recorded sites from both animals are shown in **Figure 3A** (Monkey L in Blue and Monkey V in red).

Neurons were mainly searched while the monkey freely (head-unrestrained) scanned the environment. Once reliable neuronal spiking was noticed, the experiment started. The response field of a neuron was mapped while the animal performed the memory-guided saccade. After determining the horizontal and vertical extent of the response field, we presented the targets (one per trial) in a 4 × 4 to 7 × 7 array (5 –10° from each other) ranging 30-80°. This allowed characterization of visual and motor response fields. We aimed at collecting approximately 10 trials for each target. Thus, for bigger response fields (hence more targets), a greater number of recorded trials were needed and vice versa. On average 343 ± 166 (mean ± SD) and 331 ± 156 trials/neuron were recorded in SEF and FEF respectively, again depending on the size of the response field. We did such recordings from > 200 SEF and FEF sites, often in conjunction with each other.

### Data Inclusion Criteria, sampling window and neuronal classification

In total, we isolated 256 SEF and 312 FEF neurons. Of these, we only analyzed task-modulated neurons with clear visual burst and/or with perisaccadic movement response. Neurons that only had post-saccadic activity (activity after the saccade onset) were excluded. Moreover, neurons that lacked significant spatial tuning were also eliminated (see ‘Testing for Spatial Tuning’ below). In the end, after applying our exclusion criteria, we were left with 68 SEF and 147 FEF spatially tuned neurons. We only included those trials where monkeys landed their gaze within the acceptance window for reward, however, from our analysis we removed gaze end points beyond ± 2° of the mean distribution.

### Intermediate spatial models used in main analysis

Our previous findings on FEF and SEF neurons have reported that response fields do not fit exactly against canonical models like Te or Ge, but actually may fit best against intermediate models between these canonical ones ^14^. From our previous studies ^12,15,16^ we found that a Te-Ge (T-G, target-to-gaze) continuum (specifically, steps along the ‘error line’ between Te to Ge) best quantified the egocentric visuomotor transformation in the FEF and SEF **(Fig. 3C1**), thus, in current analysis we particularly focused on this continuum. Essentially, the continuum represents a concept that is similar to an intermediate reference frame (e.g., between the eye and head) but here it is intermediate between the target and the final gaze position within the same frame of reference.

### Fitting Neural Response Fields against Spatial Models

To differentiate/test between different spatial models, conceptually, they should be spatially separable ^12,28^. The variation in natural behavior of monkeys allowed this spatial separation (see Results for details). For example, the variability produced by memory-guided gaze shifts allowed us to dissociate target coding from the gaze coding; the initial location of eye and head permitted us to differentiate between different egocentric reference frames and variability of eye and head movements for a gaze shift allowed us to distinguish different effectors. Notably, as in decoding methods that mostly test if a spatial property is implicitly coded in patterns of neuronal population activity ^70,71^, our method directly tests which model best predicts the activity in the spatially neurons. The logic of our response field fitting method is shown in **Fig. 3C2**. Specifically, if the response field activity is plotted in the correct best/reference frame, this will lead to the lowest residuals (errors between the fit and data points) in comparison with other models, i.e., if a fit calculate to its response field matches the data, then this will lead to low residuals **(Fig. 3C2**, left). Conversely, if the fit does not describe the data well, this will yield higher residuals **(Fig. 3C2**, right). For instance, an eye-fixed response field calculated in eye-coordinates will lead to lower residuals and if it is computed in any other inferior/incorrect coordinate, this will yield higher residuals^12,16.^

In reality, a non-parametric fitting method was employed to characterize the neural activity with reference to a spatial location and we also varied the kernel bandwidth of the fit to plot response field of any size, shape, or contour ^28^. The Predicted Residual Error Some of Squares (PRESS) statistics was used to test between various spatial models. To independently calculate the residual for a single trial, the actual activity associated with it was subtracted from the corresponding point on the fit calculated over all the other trials (similar to cross-validation). Importantly, if the physical shift (spatial) between two models leads to a systematic shift (direction and amount), this will be visible as a shifted or expanded response field and our model fitting method would fail to distinguish these two models as they would virtually yield indistinguishable/similar residuals. Because in our investigation, the distribution of relative positions in different models also includes a non-systematic variable component (e.g., variability in gaze endpoint errors, or pseudo-random landmark shifts), the response fields invariably were fixed at the same location, but the separation between different spatial models was based on the residual analysis.

Because the size and shape of response fields were not known beforehand and since the spatial distribution of datapoints was different for every spatial model (e.g., the models would have a higher range for eye than the head models), we calculated the non-parametric fits with different kernel bandwidths for each neuron (2-25°) thus ensuring that we did not bias the spatial fits toward a particular size and spatial distribution.

### Testing for Spatial Tuning

The model fitting method assumes that neuronal activity is structured as spatially tuned response fields, but this does suggest that other neurons do not participate in the overall population code ^72–76^ but with our analytical tool-box only tuned neurons can be explicitly tested. The neuronal spatial tuning was tested as follows. The firing rate data points were randomly (100 times to obtain random 100 response fields) shuffled across the position data that we got from the best model. We then statistically compared the mean PRESS residual distribution (PRESS_random_) of the 100 randomly generated response fields with the mean PRESS residual (PRESS_best-fit_) distribution of the best-fit model (unshuffled, original data). If the best-fit mean PRESS was outside the 95% confidence interval of the distribution of the shuffled mean PRESS, we then deemed the neuron’s activity as selective. At the spatiotemporal level, some neurons were spatially tuned at certain time-steps and others were untuned because of low signal/noise ratio. We thus removed the time steps where the populational mean spatial coherence (goodness of fit) was statistically indistinguishable from the baseline (before target onset) since there was no task-related information at this time and thus neural activity had no spatial tuning. We defined an index (Coherence Index, CI) for spatial tuning of a single neuron which was calculated as ^12^:

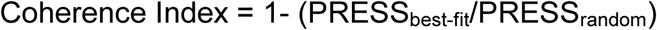

If the PRESS_best-fit_ was similar to PRESS_random_ then the CI would be roughly 0, whereas if the best-fit model is a perfect fit (i.e., PRESS_best-fit_ = 0), then the CI would be 1. We only included those neurons in our analysis that showed significant spatial tuning.

### Spatiotemporal analysis

A major goal of this study was to track the progression of the T-G code in spatially tuned neuron populations, from the visual response onset until the mask offset / landmark shift. To finely track the evolution of the spatiotemporal code, we smoothed and binned the activity form visual response onset until the landmark shift into 7 half-overlapping bins. To this aim, the neural firing rate (in spikes/second; the number of spikes divided by the sampling interval for each trial) was sampled into 7 half-overlapping time windows (with a width of 120 ms). The bin number was chosen in such a way so that the sampling time window was wide enough, and thus robust enough to account for the stochastic nature of neuronal spiking activity (ensuring that there were enough neuronal spikes in the sampling window for effective spatial analysis) ^13,16^. Once we estimated the firing rate for each trial at a given time-step, they were pooled together for spatial modeling. This procedure allowed us to treat the whole sequence of visual-memory responses from the visual response onset until the onset of landmark shift as a continuum.

## Acknowledgement

This project was supported by a Canadian Institutes for Health Research (CIHR) Grant and the Vision: Science to Applications (VISTA) Program, which is supported in part by the Canada first Research Excellence Fund and by Deutsche Forschungsgemeinschaft (IRTG-1901, RU-1847 and CRC/TRR-135, project number 222641018). VB, XY, and HW are supported by CIHR and VISTA. PAK is supported by VISTA. JDC is supported by the Canada Research Chair Program.

## Supplementary Figures

**Supplementary Figure 1:**
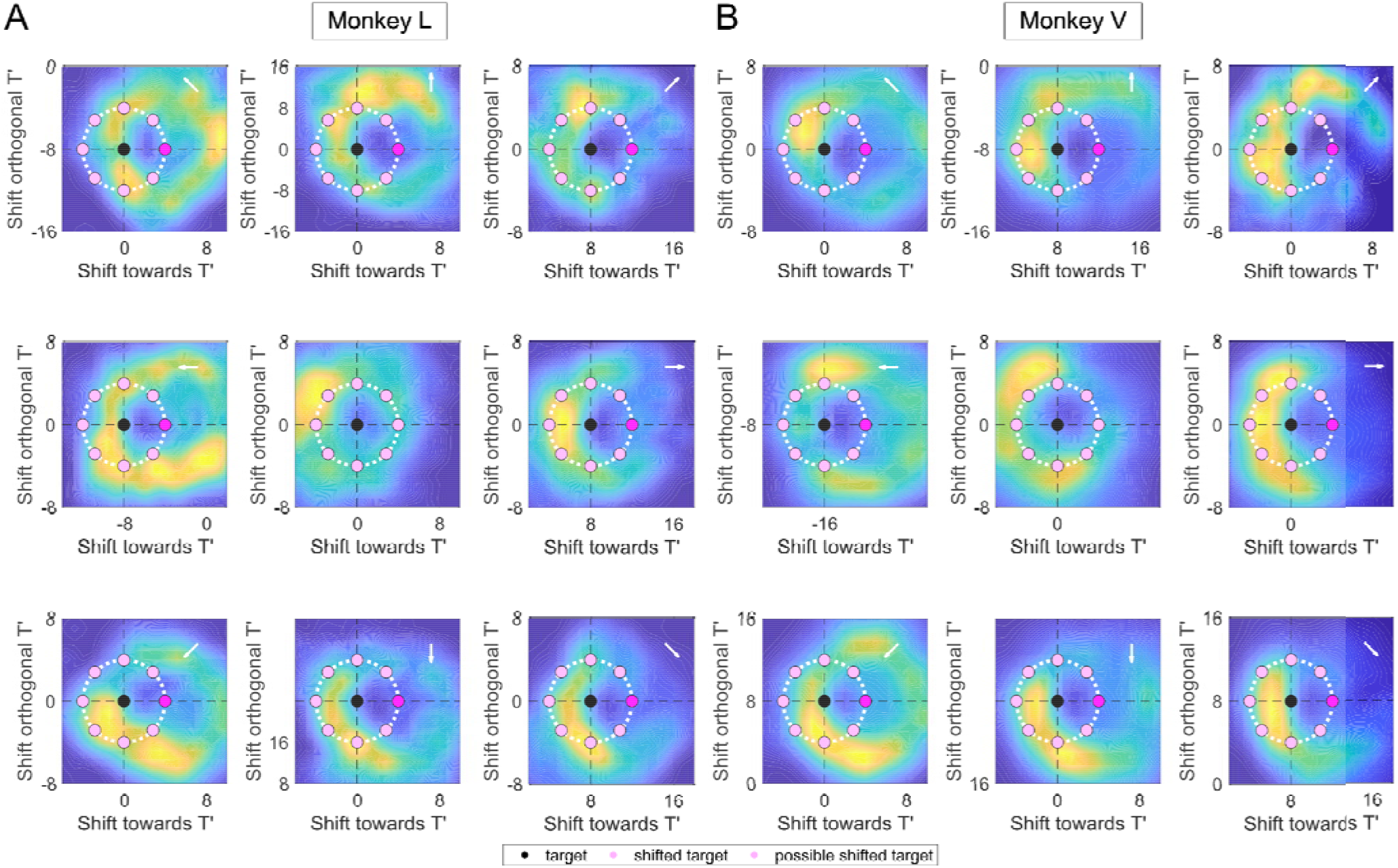
The ‘donut like’ pattern was also observed when the data were analyzed separately for each of the eight individual shift directions for both animals (**A**: Monkey L; **B**: Monkey V). The white arrow indicates the direction of the shift.

**Supplementary Figure 2:**
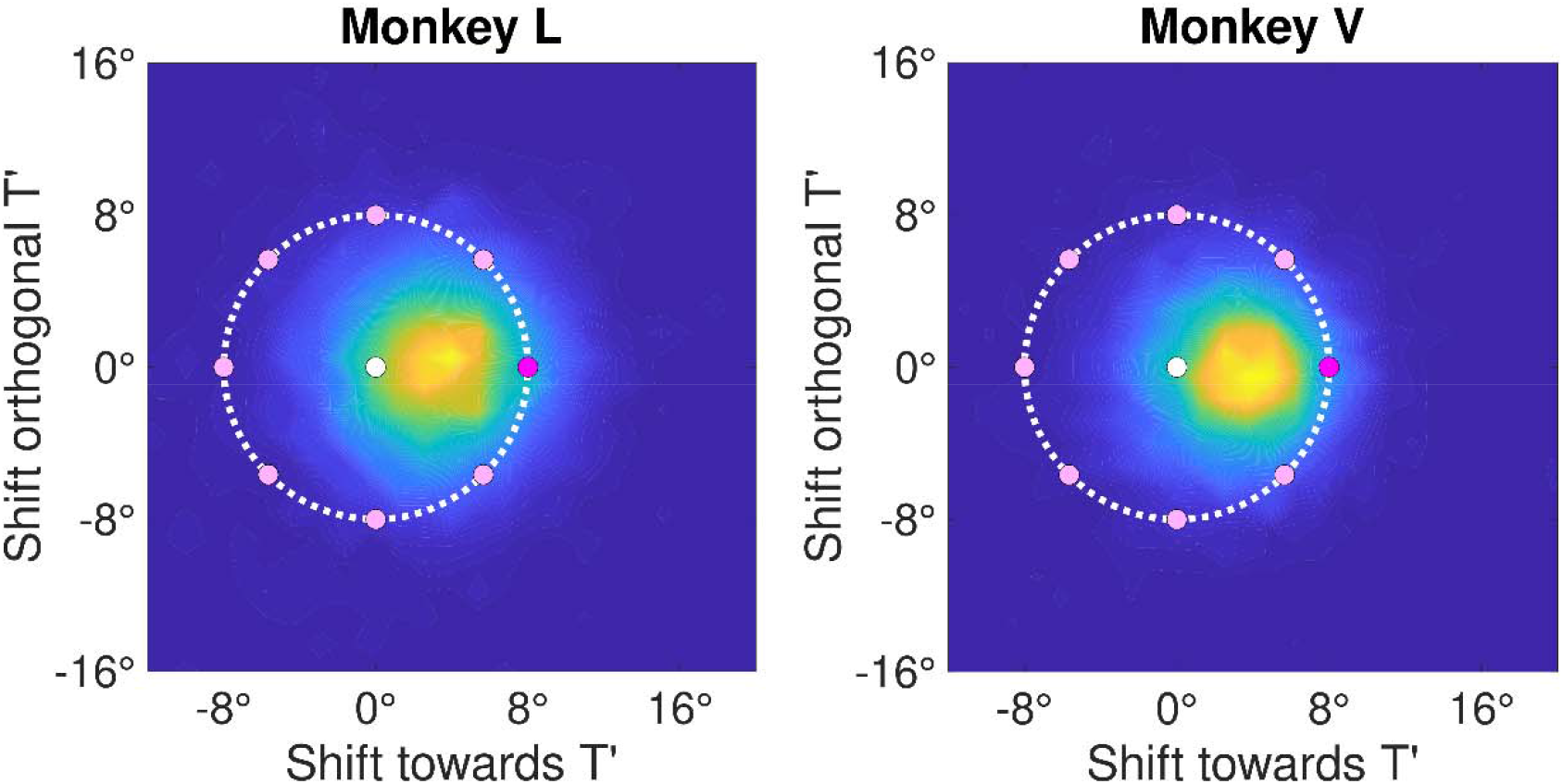
Subtraction of the no-shift trials from the shift trials leads to the collapse of torus into a gaussian distribution in both animals (Left: Monkey L, Right: Monkey V).

